# The *Bacillus subtilis* class A penicillin-binding protein 4 (PBP4) requires an accessory protein RpdA

**DOI:** 10.64898/2025.12.21.695826

**Authors:** Ruoqi Huang, Yesha Patel, John D. Helmann

## Abstract

Peptidoglycan (PG) synthesis is essential to maintain cell integrity during bacterial growth and division. In *Bacillus subtilis,* PG synthesis involves class A PBPs that act independently and class B PBPs that function in complexes for cell elongation (the elongasome) and division (the divisome). Here, we identify RpdA (formerly YufK) as a specific regulator of PBP4. Inactivation of either vegetative class A PBP (PBP1 or PBP4) by the β-lactam antibiotic cefuroxime (CEF) is toxic and their loss confers CEF resistance. Similarly, loss of RpdA increases CEF resistance and genetic epistasis studies reveal that RpdA functions in a pathway with PBP4. In the absence of RpdA, PBP4 is no longer membrane localized. Analysis of a predicted RpdA-PBP4 protein complex suggests that RpdA has a second function in addition to serving as a membrane scaffold. Induction of RpdA reduces sensitivity to an undecaprenyl-phosphate binding antibiotic, consistent with a role in recycling of this important lipid carrier. We conclude that RpdA is a PBP4 accessory protein critical for its localization and activity.

## Introduction

Most bacterial cells possess a rigid, mesh-like cell wall that surrounds their cytoplasmic membrane. This cell wall is primarily composed of peptidoglycan (PG), a complex polymer consisting of linear chains of disaccharides N-acetyl glucosamine (NAG) and N-acetylmuramic acid (NAM) cross-linked via short pentapeptide stems (Anderson *et al*, 2025; Vollmer Blanot & de Pedro, 2008). The PG layer provides mechanical strength, maintains cellular morphology, and protects bacterial cells from osmotic lysis. In most Gram-negative bacteria, PG is relatively thin and lies beneath an outer membrane enriched in lipopolysaccharides (Deghelt Pierre Despas & Collet, 2025). In contrast, Gram-positive bacteria lack this outer membrane but compensate with a considerably thicker PG layer. PG biosynthesis is essential for bacterial viability, and most clinically used antibiotics act by inhibiting this process (Mora-Ochomogo & Lohans, 2021; Ntallis *et al*, 2025).

PG formation is initiated in the cytoplasm with the synthesis of its monomeric sugar precursors, NAG and NAM (Barreteau *et al*, 2008). Together with a pentapeptide side-chain, these precursors are linked to undecaprenyl pyrophosphate (UPP) to form lipid II, which is then flipped across the membrane (Kumar *et al*, 2022). Subsequent polymerization and incorporation into the cell wall are driven by penicillin-binding proteins (PBPs) (Sauvage *et al*, 2008). Class A PBPs (aPBPs) can independently catalyze both transglycosylation (TG) and transpeptidation (TP) reactions, whereas class B PBPs ((Paradis-Bleau *et al*, 2010)s) catalyze only the TP reaction and depend on SEDS (Shape, Elongation, Division, and Sporulation) family proteins to provide the TG function (Galinier *et al*, 2023; Terrak & Kerff, 2025). In the model Gram-positive bacterium *Bacillus subtilis,* the elongasome is essential and includes the SEDS protein RodA (TG) and either of two class B PBPs (TP) (Emami *et al*, 2017; Meeske *et al*, 2016). In contrast, all four aPBPs are individually and collectively dispensable under standard growth conditions (McPherson & Popham, 2003). However, mutants lacking all aPBPs display altered cell morphology and impaired growth, highlighting the physiological significance of these enzymes (Dion *et al*, 2019).

The enzymes involved in PG synthesis are prime targets for antibiotics (Ntallis *et al*., 2025). β-lactams are broad spectrum antibiotics that bind and inhibit PBPs by covalently binding to their TP domains (Mora-Ochomogo & Lohans, 2021). Cefuroxime (CEF) preferentially targets aPBPs (Sharifzadeh *et al*, 2020) and disrupts cross-linking of linear glycan strands resulting in the accumulation and subsequent degradation of uncrosslinked PG, a process known as futile cycling (Cho Uehara & Bernhardt, 2014). This energetically wasteful cycle contributes to β-lactam-induced lysis. However, in mutants lacking PBP1 (*ponA*) or PBP4 (*pbpD*) futile cycling is reduced and cells have reduced CEF sensitivity.

*B. subtilis* employs multiple regulatory systems for PG synthesis (Galinier *et al*., 2023; Helmann, 2016). At the transcriptional level, the antibiotic-induced extracytoplasmic function (ECF) sigma factor σ^M^ controls over 200 genes (Eiamphungporn & Helmann, 2008). These genes encode proteins for the synthesis and flipping of the lipid II precursor, for PG polymerization, and for dephosphorylation of UPP (BcrC) and flipping of the resulting UndP (UptA) (Roney & Rudner, 2023). Another stress-induced sigma factor, σ^I^, upregulates the elongasome components MreBH and LytE (Patel Zhao & Helmann, 2020). Additionally, the WalRK two-component system regulates expression of the autolysins LytE and CwlO (Dobihal *et al*, 2022). In addition to transcriptional control, PG synthesis is also regulated at the level of protein–protein interactions (Egan Errington & Vollmer, 2020; Galinier *et al*., 2023). Elongasome activity is regulated by RagB, which binds to RodA (Pompeo *et al*, 2025), and TseB, which interacts with PBP2A (Delisle *et al*, 2021). Activity of the major class A PBP, PBP1, is regulated by GpsB-dependent complex assembly (Cleverley *et al*, 2019) and by an extracytoplasmic intrinsically disordered region (IDR) that detects gaps in the PG meshwork (Brunet *et al*, 2022).

In this study, we identify and characterize a **<u>r</u>**egulator of **<u>P</u>**BP4 (*pb<u>p**D**</u>*) **<u>a</u>**ctivity, RpdA (formerly YufK). A Δ*rpdA* strain exhibited reduced susceptibility to CEF, similar to strains lacking PBP1 or PBP4. This resistance was additive with Δ*ponA* but not with Δ*pbpD*, suggesting that RpdA specifically affects PBP4 activity. Cell fractionation and structural modeling suggest that RpdA functions as a scaffold for anchoring PBP4 on the membrane. Intriguingly, induction of RpdA reduces the sensitivity of a Δ*uptA* strain (lacking the major UndP transporter) to MX2401, an antibiotic that binds UndP. Together, these findings reveal a previously uncharacterized accessory protein that regulates aPBP localization and activity and may additionally serve as an UndP transporter.

## Results

### An *rpdA* deletion reduces cefuroxime sensitivity

In a previous study, we identified a strain carrying a mutation (H482Y) in the RNA polymerase β-subunit to be significantly more sensitive to cefuroxime (CEF), a β-lactam antibiotic (Patel & Helmann, 2025; Patel *et al*, 2023). To uncover genetic changes that can confer CEF resistance in this background (RpoB^H482Y^), we screened for suppressor mutations and through whole genome sequencing identified alterations in *rpdA* (formerly *yufK*) (SI Table 1). Both mutations were loss-of-function alleles. Increased CEF resistance was confirmed by disk diffusion assay, where Δ*rpdA* in the RpoB^H482Y^ background led to a reduced zone of inhibition compared to the parent strain, as also observed for the original suppressors (SI Fig 1).

**Fig 1.**
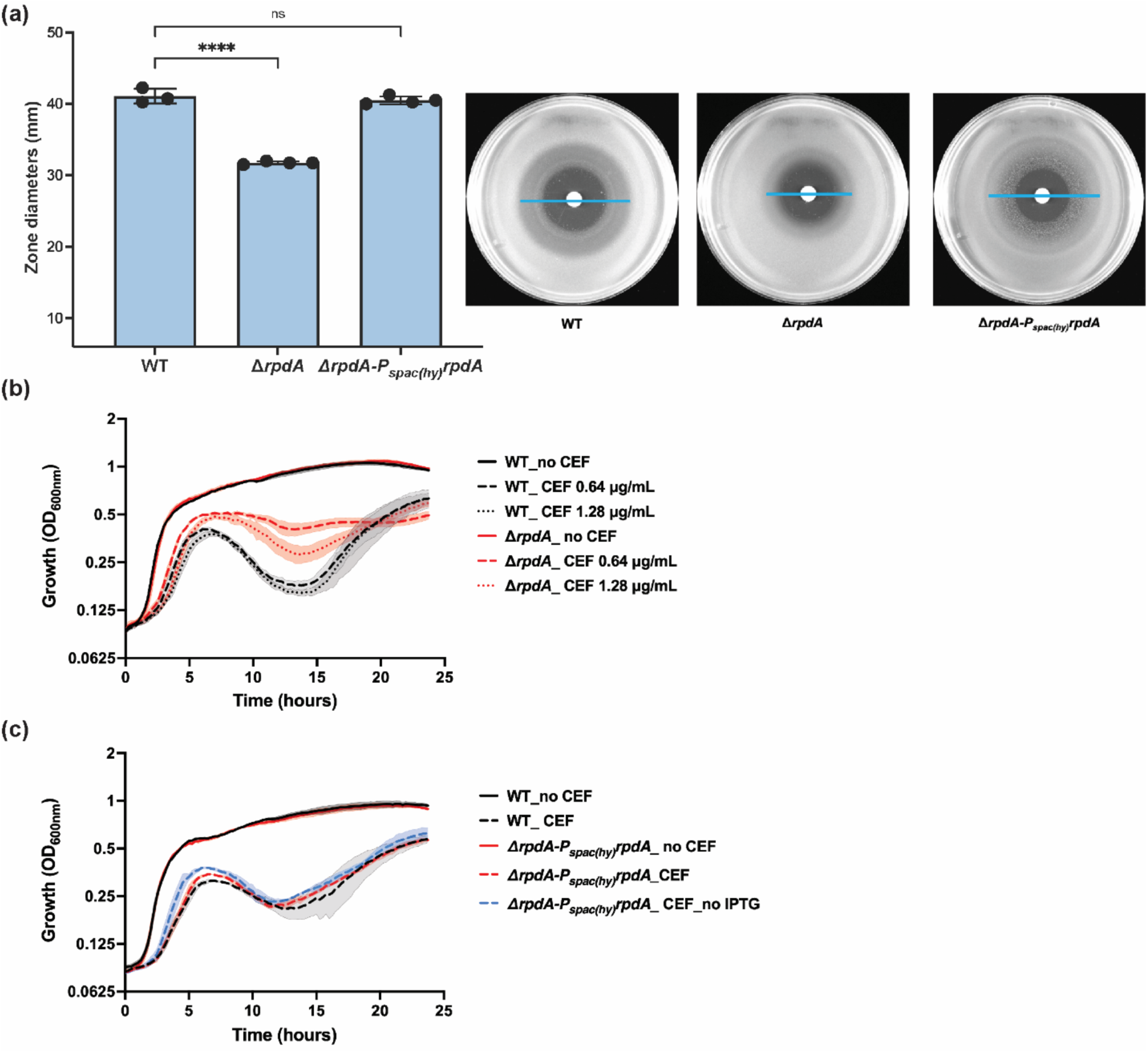
An *rpdA* deletion reduces cefuroxime (CEF) sensitivity. **(a)** Disk diffusion assay of WT, Δ*rpdA* and Δ*rpdA* complemented with *rpdA* (Δ*rpdA amyE*::P*_spac(hy)_-rpdA*) using disks loaded with 30 μg CEF. Plates were supplemented with 0.5 mM IPTG. The diameters of the zone of inhibition include the zone of no growth and poor growth. Representative images for each strain are shown, with the zone of inhibition indicated by a blue line. The y-axis begins at 6 mm, corresponding to the disc diameter, which was subtracted from all measurements. Data are represented as mean ± standard deviation (SD) for at least three biological replicates. Statistical significance was determined by one-way ANOVA with Dunnett’s correction; **** indicate *p* < 0.0001; ns indicate not significant. **(b)** Growth kinetics of WT (black) and Δ*rpdA* (red) strains in the absence (solid lines) or presence of 0.64 μg/mL (dashed lines) and 1.28 μg/mL (dotted lines) CEF. Data are represented as mean from six biological replicates; shaded regions indicate SD. **(c)** Growth kinetics of WT (black) and Δ*rpdA* complemented with *rpdA* (Δ*rpdA amyE*::P*spac(hy)-rpdA*). Solid lines indicate growth without CEF, and dashed lines represent growth in the presence of 0.64 μg/mL CEF. Complementation was apparent in both the presence (red) or absence (blue) of 0.5 mM IPTG. Data are represented as mean from at least three biological replicates; shaded regions indicate SD.

**Table 1.** Minimum inhibitory concentrations (MICs) of cefuroxime (CEF) for WT and Δ*rpdA* strains. MICs were determined using E-test strips on lysogeny broth (LB) medium and Mueller–Hinton (MH) agar plates. Values represent mean ± standard deviation (SD) from three biological replicates.

|  | Medium | MIC (μg/mL) |
| --- | --- | --- |
| WT | LB | 3.50 ± 0.84 |
|  | MH | 3.67 ± 0.52 |
| *ΔrpdA* | LB | 8.33 ± 1.97 |
|  | MH | 10.00 ± 2.19 |

To determine whether Δ*rpdA* reduces CEF sensitivity independent of the *rpoB* mutation, we deleted *rpdA* in the wild-type 168 (WT) strain. Compared to WT, the Δ*rpdA* strain had a smaller zone of inhibition, indicating reduced sensitivity to CEF (Fig. 1a). Consistently, the minimum inhibitory concentration (MIC) for CEF was two-fold higher for Δ*rpdA* compared to WT (Table 1).

In liquid media, Δ*rpdA* grew like WT in the absence of CEF. After 5 hours of treatment with 0.64 or 1.28 μg/mL CEF, WT cells exhibited a reduction in OD_600_, likely reflecting cell lysis, followed by recovery of growth after 15 hours. In contrast, Δ*rpdA* strains had significantly reduced cell lysis (Fig. 1b). To determine whether the drop in optical density correlated with a loss of viable cell counts, we measured colony forming units (CFUs) at 12, 15 and 18 hours post treatment with 1.28 μg/mL CEF. In WT cells, the two-fold drop in OD_600_ corresponded to a ten-fold reduction in CFU, confirming that the decrease in optical density reflects a loss of viability. The smaller change in OD_600_ compared to CFUs may result from non-viable but unlysed cells. In contrast, the Δ*rpdA* mutant showed no significant reduction in CFUs (SI Fig. 2). Complementation of Δ*rpdA* by ectopic induction of *rpdA* with 0.5 mM IPTG restored CEF sensitivity to WT levels (Fig. 1c), confirming that *rpdA* deletion contributed to CEF resistance. The complementation was IPTG-independent due to the leakiness of the P*_spac(hy)_* promoter used here (Castillo-Hair *et al*, 2019).

**Fig 2.**
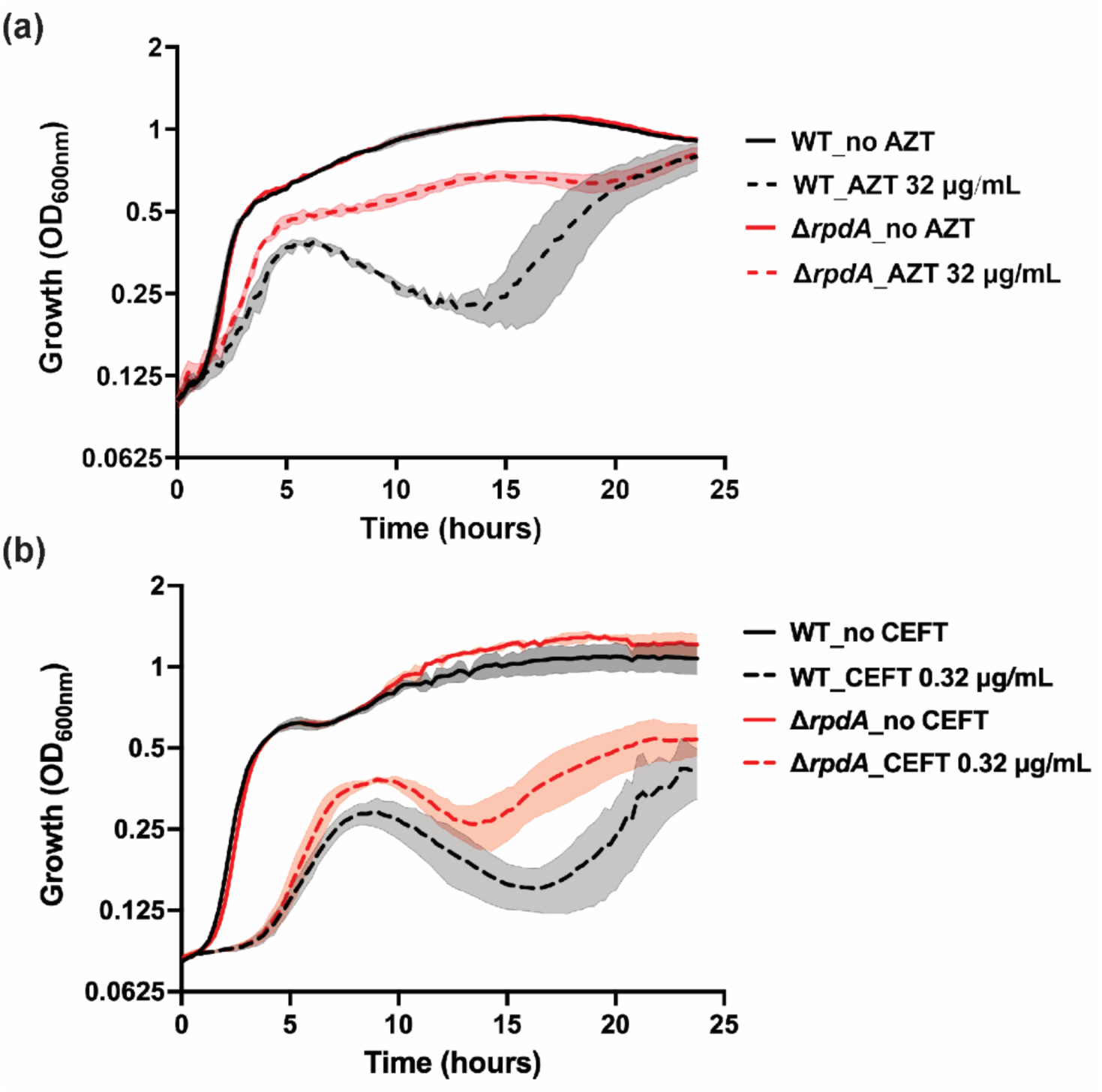
An *rpdA* deletion reduces sensitivity to β-lactams targeting aPBPs. Growth kinetics of WT (black) and Δ*rpdA* (red) strains in the presence (dashed lines) or absence (solid lines) of **(a)** aztreonam (AZT, 32 μg/mL) and **(b)** ceftriaxone (CEFT, 0.32 μg/mL). Data are represented as mean from at least three biological replicates; shaded regions indicate SD.

### An *rpdA* deletion specifically reduces sensitivity to aPBP-targeting β-lactams

Next, we determined the effect of *rpdA* deletion against a panel of cell wall-targeting antibiotics. These antibiotics included different β-lactams (aztreonam, ceftriaxone, penicillin G, methicillin, cephalexin, oxacillin, cloxacillin, mecillinam), as well as non-β-lactam inhibitors of cell wall synthesis (moenomycin, bacitracin, vancomycin, fosfomycin). Growth (OD_600_) of WT and Δ*rpdA* was monitored on treatment with all antibiotics except cloxacillin and mecillinam, for which MICs were determined using E-test strips. Among all the antibiotics tested, Δ*rpdA* cells exhibited significantly reduced lysis upon treatment with antibiotics previously shown to preferentially target aPBPs (Sharifzadeh *et al*., 2020). Specifically, Δ*rpdA* reduced susceptibility to 32 μg/mL of aztreonam (AZT; Fig. 2A), 0.32 μg/mL of ceftriaxone (CEFT; Fig. 2B), and 0.08 and 0.16 μg/mL of cephalexin (SI Fig 3). For all other antibiotics, including other β-lactams, inactivation of *rpdA* did not reduce susceptibility (SI Fig. 3). Together, these findings suggest that RpdA function may specifically be linked to aPBP activity.

**Fig 3.**
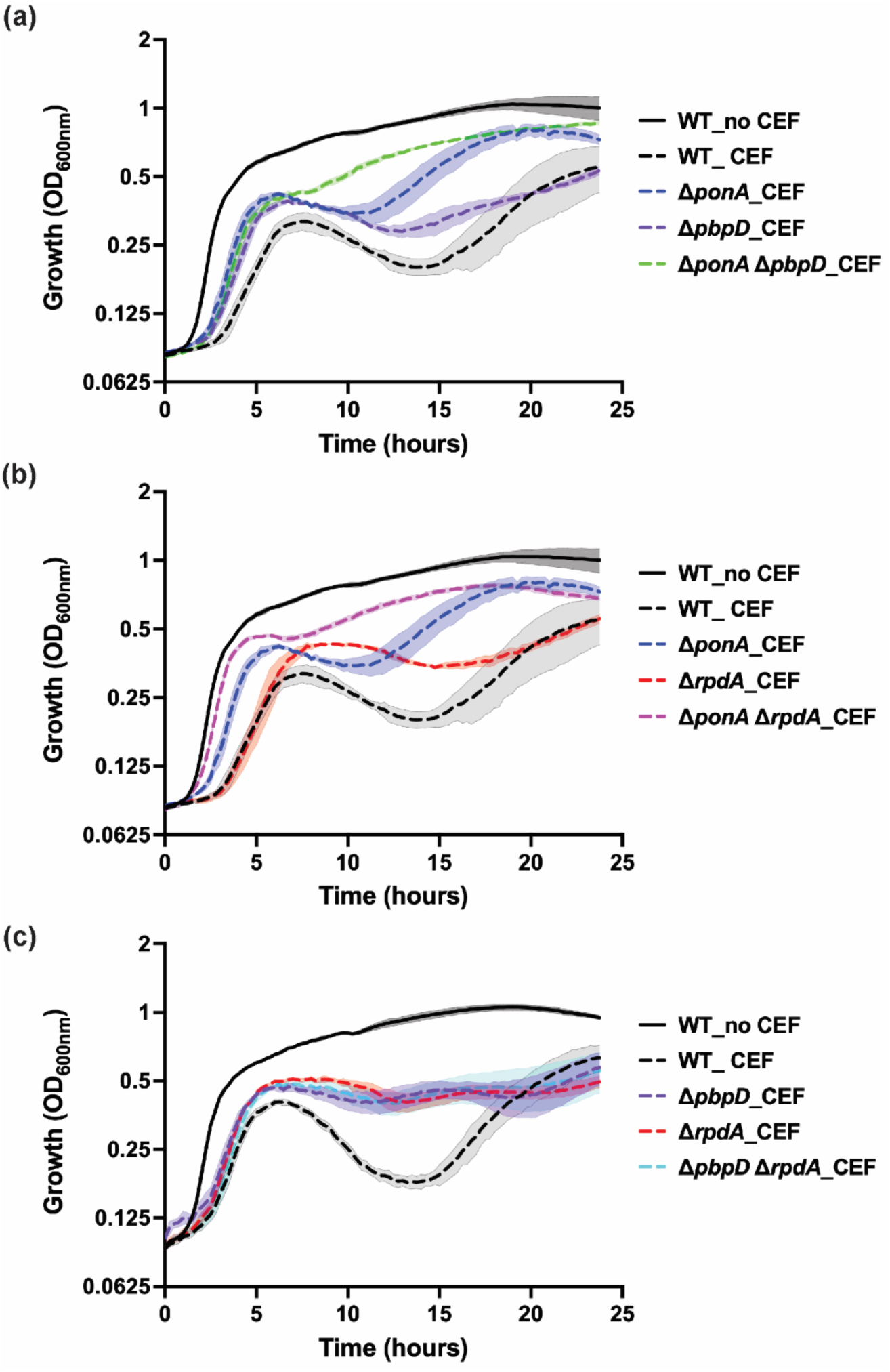
*rpdA* and *pbpD* exhibit epistasis. Growth kinetics of WT (black) and **(a)** Δ*pbpD* (purple), Δ*ponA* (blue) and Δ*ponA* Δ*pbpD* double mutant (green). **(b)** Δ*rpdA* (red), Δ*ponA* (blue) and Δ*ponA* Δ*rpdA* double mutant (magenta) **(c)** Δ*pbpD* (purple), Δ*rpdA* (red) and Δ*pbpD* Δ*rpdA* double mutant (cyan) in the presence (dashed lines) or absence (solid lines) of 0.64 μg/mL CEF. Data are represented as mean from at least three biological replicates; shaded regions indicate SD.

### RpdA is epistatic with PBP4

In *B. subtilis*, PBP1 (encoded by *ponA*) and PBP4 (*pbpD*) are aPBPs expressed during vegetative growth, whereas PBP2c (*pbpF*) and PBP2d (*pbpG*) are primarily expressed during sporulation (McPherson Driks & Popham, 2001; Straume *et al*, 2021). To determine which of these aPBPs contribute to CEF sensitivity, we tested the response of single deletion mutants to 0.64 μg/mL CEF. Strains lacking either *ponA* or *pbpD* had significantly reduced lysis in response to CEF treatment, similar to the Δ*rpdA* mutant (Fig. 3a). In contrast, deletion of *pbpF* and *pbpG* had no effect on the CEF-induced cell lysis (SI Fig. 4a), consistent with their sporulation-specific expression. Moreover, the Δ*ponA* Δ*pbpD* double mutant showed reduced lysis compared to either single mutant, suggesting that PBP1 and PBP4 have additive roles in mediating CEF sensitivity (Fig. 3a).

**Fig 4.**
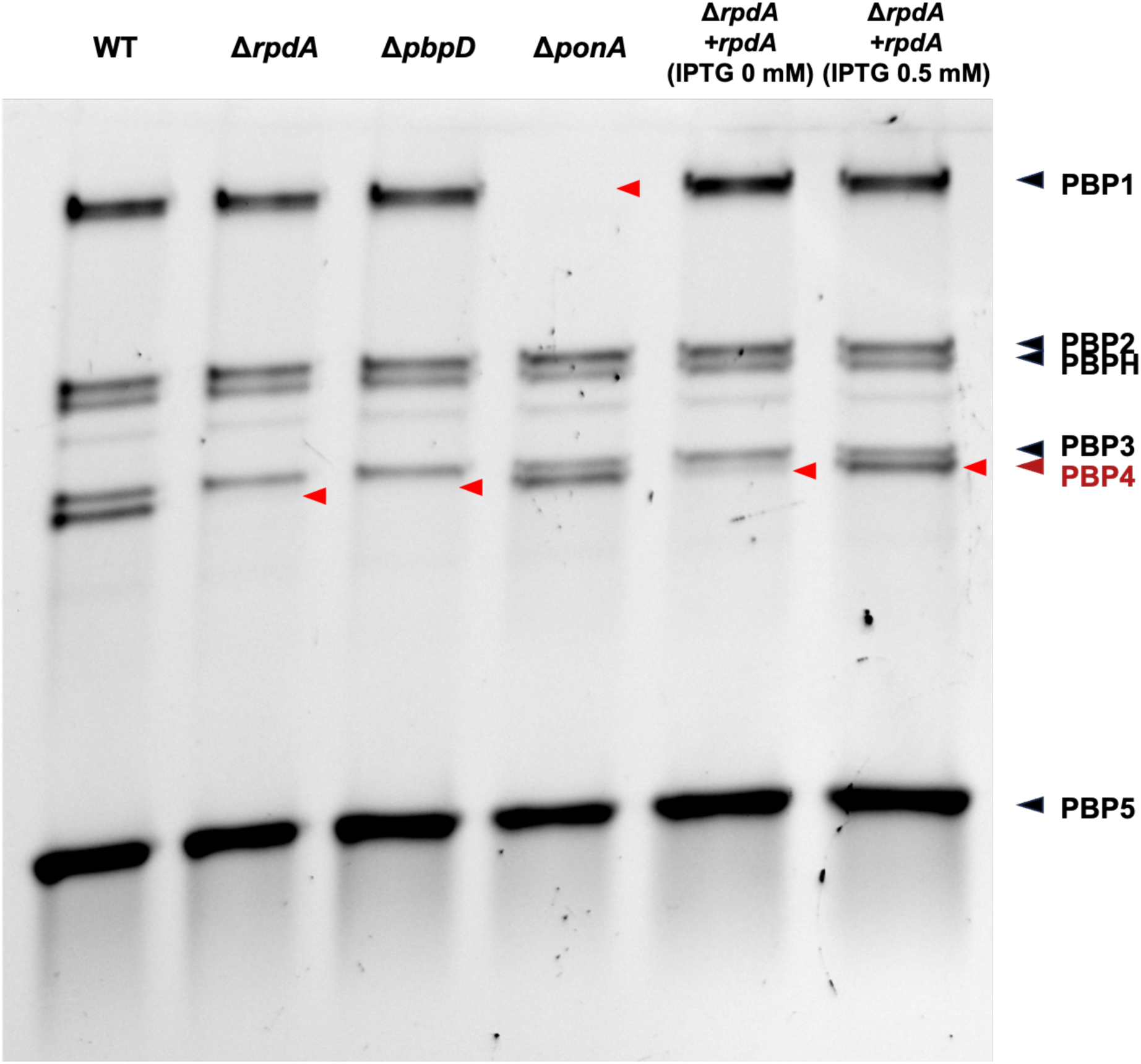
*R*pdA is essential for PBP4 transpeptidase activity. Bocillin-FL activity-based probe assay was performed using membrane fractions from WT, Δ*rpdA*, Δ*pbpD*, Δ*ponA* and Δ*rpdA amyE*::P*_spac(hy)_-rpdA* strains (indicated as Δ*rpdA* + *rpdA*) grown in the presence and absence of 0.5 mM IPTG. Labeled proteins were separated by SDS-PAGE and visualized by fluorescence scanning. Red arrows on the gel indicate the loss of PBP4 band in the Δ*rpdA* and Δ*pbpD* strain and the PBP1 band in Δ*ponA* strain. The experiment was independently repeated three times; a representative image from one biological replicate is shown. This is a cropped gel image. A full image with other mutant strains is in SI Fig. 6.

The similar phenotypes observed in Δ*ponA*, Δ*pbpD*, and Δ*rpdA* mutants suggested that RpdA may impact the activity of PBP1, PBP4, or both. To test if Δ*rpdA* is epistatic with Δ*ponA* or Δ*pbpD*, we constructed Δ*ponA* Δ*rpdA* and Δ*pbpD* Δ*rpdA* double mutants. On treatment with 0.64 μg/mL CEF, Δ*ponA* Δ*rpdA* had improved growth with a shorter lag phase and reduced lysis, compared to both the single mutants (Fig. 3b). In contrast, Δ*pbpD*, Δ*rpdA*, and the Δ*pbpD* Δ*rpdA* double mutant had identical responses on CEF treatment (Fig. 3c). We also deleted *rpdA* in a Δ*ponA* Δ*pbpD* double mutant and a Δ4 strain missing all the 4 aPBPs (SI Fig. 4b) In every case, Δ*rpdA* had no additional effect in any strain lacking *pbpD*, consistent with the hypothesis that RpdA and PBP4 act in the same pathway. We considered the possibility that RpdA might regulate *pbpD* expression. However, using real-time PCR we found that *pbpD* transcript levels were only slightly reduced (<2-fold) by *rpdA* deletion (SI Fig. 5).

**Fig 5.**
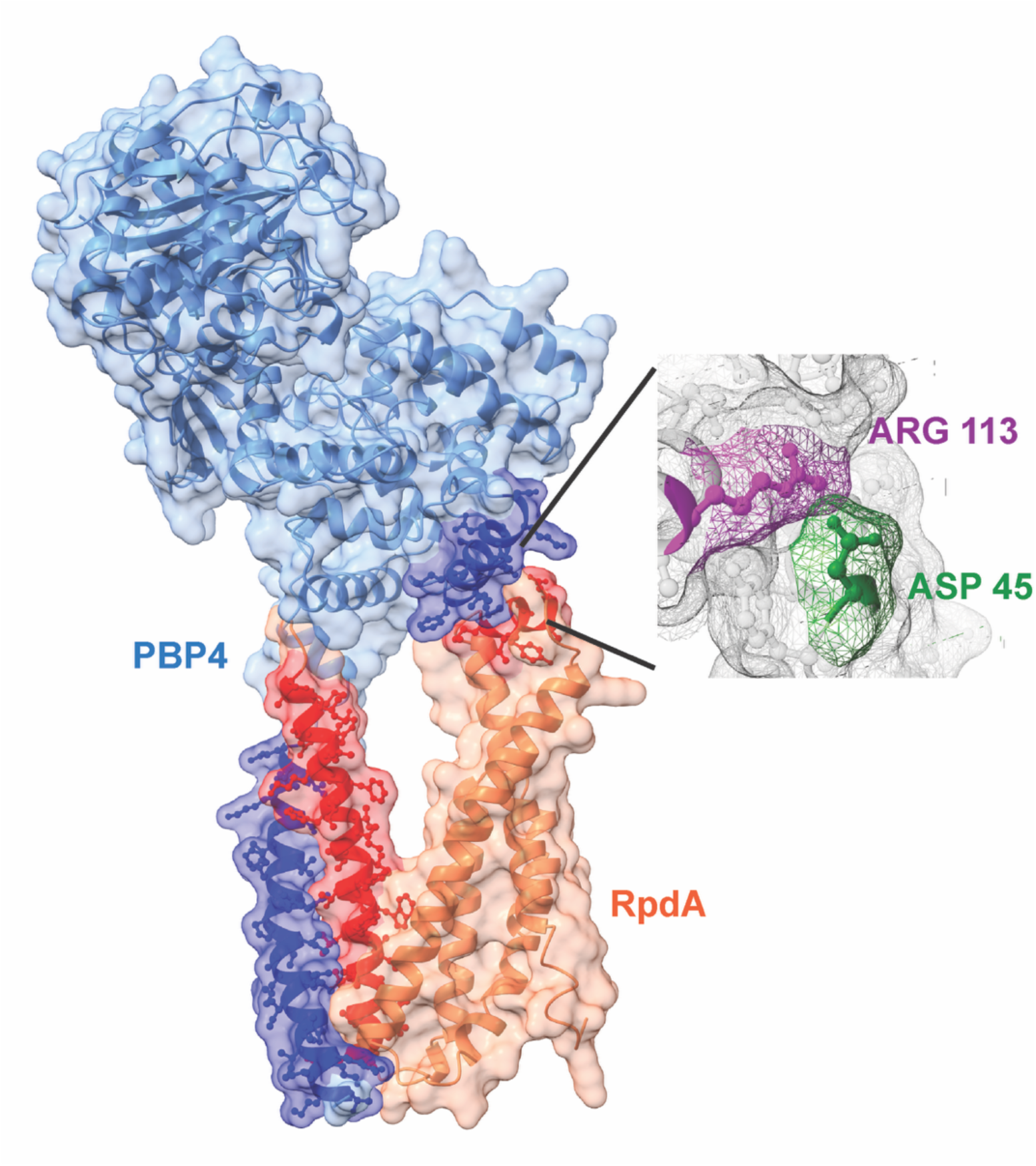
Predicted structure of the PBP4–RpdA complex. AlphaFold v3 prediction of the PBP4 (blue)–RpdA (coral) complex (ipTM = 0.77) (Abramson *et al*., 2024). The dark blue and dark coral α-helices represent the regions predicted to interact, potentially mediating membrane tethering of PBP4 or other functions affecting the PBP4 activity.

### PBP4 activity was lost in the absence of RpdA

To determine the effect of loss of RpdA on PBP4 activity, we used bocillin-FL (Boc-FL), a fluorescent penicillin V analog that binds covalently to the transpeptidase domains of PBPs (Sharifzadeh *et al*., 2020). When PBPs are properly folded and catalytically active, Boc-FL irreversibly labels the serine residue in the active site, enabling visualization of individual PBPs as distinct bands on SDS-PAGE. Loss or reduction of a specific band indicates decreased Boc-FL binding, which may result from reduced protein abundance or impaired transpeptidase activity of the PBP. We treated WT, Δ*rpdA*, Δ*pbpD*, Δ*ponA* mutants and Δ*rpdA* strains carrying an IPTG-inducible copy of *rpdA* (with and without 0.5 mM IPTG) with Boc-Fl (Fig. 4). As expected, the membrane fractions of both Δ*pbpD* and Δ*ponA* lacked PBP4 and PBP1 bands respectively. Strikingly, the Δ*rpdA* mutant (and Δ*ponA* Δ*rpdA* double mutant; SI Fig. 6) also lacked the PBP4 band, suggesting that PBP4 is inactive or undetectable in the absence of RpdA. In the Δ*rpdA* strain harboring the inducible *rpdA* construct, leaky (uninduced) expression resulted in a faint PBP4 band, while induction with 0.5 mM IPTG restored a strong PBP4 signal (Fig. 4). Together, these results confirmed that RpdA was required for PBP4 activity. Importantly, no appreciable changes were observed in the levels of other PBPs in the Δ*rpdA* strain compared to WT, indicating PBP4 activity depends on RpdA.

### A predicted physical interaction between RpdA and PBP4

RpdA is an uncharacterized membrane protein with a DUF5366 domain and predicted to contain five transmembrane helices (Fig. 5, coral subunit). Consistent with the hypothesis that RpdA may physically interact with PBP4 (Fig. 5, blue subunit), AlphaFold v3 predicts a RpdA–PBP4 complex with an ipTM value of 0.77 (Fig. 5). In the predicted complex, one of the transmembrane α-helices of RpdA (dark coral) closely associates with the membrane-spanning α-helical domain of PBP4 (dark blue), suggesting that RpdA may function as a scaffold that anchors PBP4 to the membrane. In addition, a probable interaction was also observed between RpdA and the transglycosylase domain of PBP4 on the extracellular side of the membrane that may also affect PBP4 activity.

### RpdA is required for membrane localization of PBP4

To test the hypothesis that RpdA may help localize PBP4 to the membrane, we examined the presence of PBP4-FLAG in whole-cell lysates and membrane fractions of WT and Δ*rpdA* strains by western blotting (Fig. 6, SI Fig. 7). PBP4-FLAG was readily detected in whole-cell lysates of the Δ*rpdA* mutant, indicating that PBP4 protein levels, like *pbpD* mRNA levels (SI Fig. 5), were not significantly altered in the absence of RpdA. Strikingly, PBP4-FLAG was undetectable in the membrane fraction of Δ*rpdA*, strongly suggesting that RpdA was required to stabilize or retain PBP4 in the membrane. In the absence of RpdA, PBP4 likely failed to localize correctly to the membrane, resulting in a loss of its functional activity. Consistent with this model, both *rpdA* mutations identified in our preliminary screen for CEF-resistance (SI Table 1) truncate the C-terminal α-helix. We speculate that these mutations also conferred CEF-resistance due to mislocalization of PBP4.

**Fig 6.**
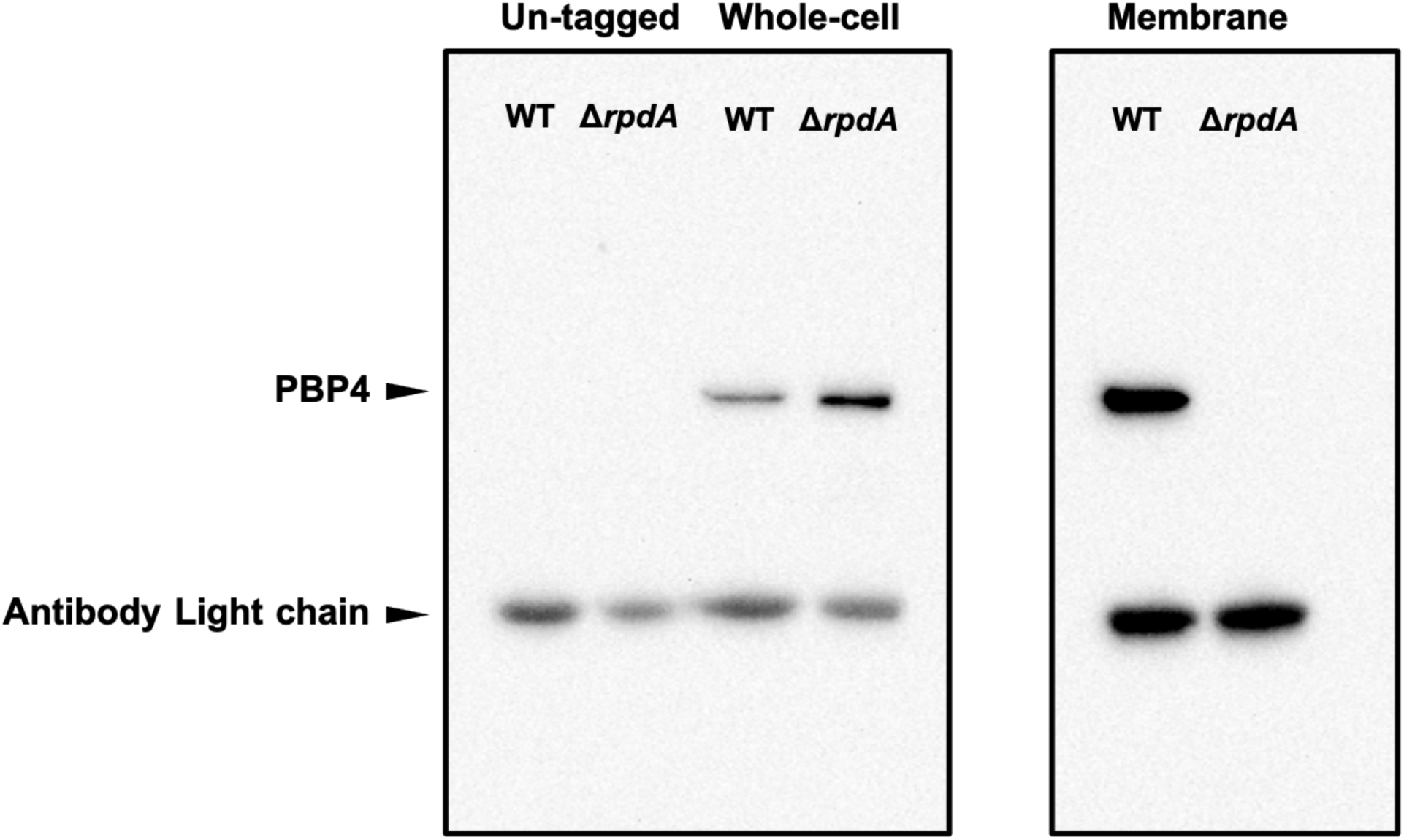
RpdA is essential for membrane localization of PBP4. Western blot analysis of PBP4-FLAG in whole-cell lysate and membrane fractions of WT and Δ*rpdA* strains. WT and Δ*rpdA* strains without FLAG tagged PBP4 were used as negative control (Untagged fraction). PBP4-FLAG was enriched using anti-FLAG magnetic beads. A consistent antibody light chain band visible in all samples is from the antibody-coated beads. The experiment was independently repeated three times; a representative image is shown. Stain-free gel imaging (SI Fig. 7a) was used to compare the amount of protein loaded in each well shown. A composite blot image with the protein ladder is shown in SI Fig. 7b.

### RpdA may have additional roles in supporting PBP4 activity

Next, we tested whether the predicted interaction between the extracellular α-helices of PBP4 and RpdA also contributes to PBP4 activity. Structural analysis indicated that D45 of RpdA may form an electrostatic bond with R113 of PBP4 (Fig. 5, inset). We generated an RpdA^D45R^ variant, with the negatively charged aspartate substituted by a positively charged arginine residue to weaken the RpdA–PBP4 association. Compared to the WT cells, the RpdA^D45R^ mutant exhibited increased resistance to CEF; however, the resistance was lower than that observed in the Δ*rpdA* mutant (Fig. 7A). A similar result was obtained on treatment with AZT (Fig. 7B). In addition, using Boc-FL we determined that the levels of active PBP4 on the membrane were not altered in the RpdA^D45R^ variant strain (Fig. 7C, 7D, SI Fig. 8), suggesting that the increased CEF resistance on disrupting the extracellular interaction was not dependent on PBP4 mislocalization. Together, these results suggest that, in addition to tethering PBP4 to the membrane, RpdA may play an additional role in supporting PBP4 activity.

**Fig 7.**
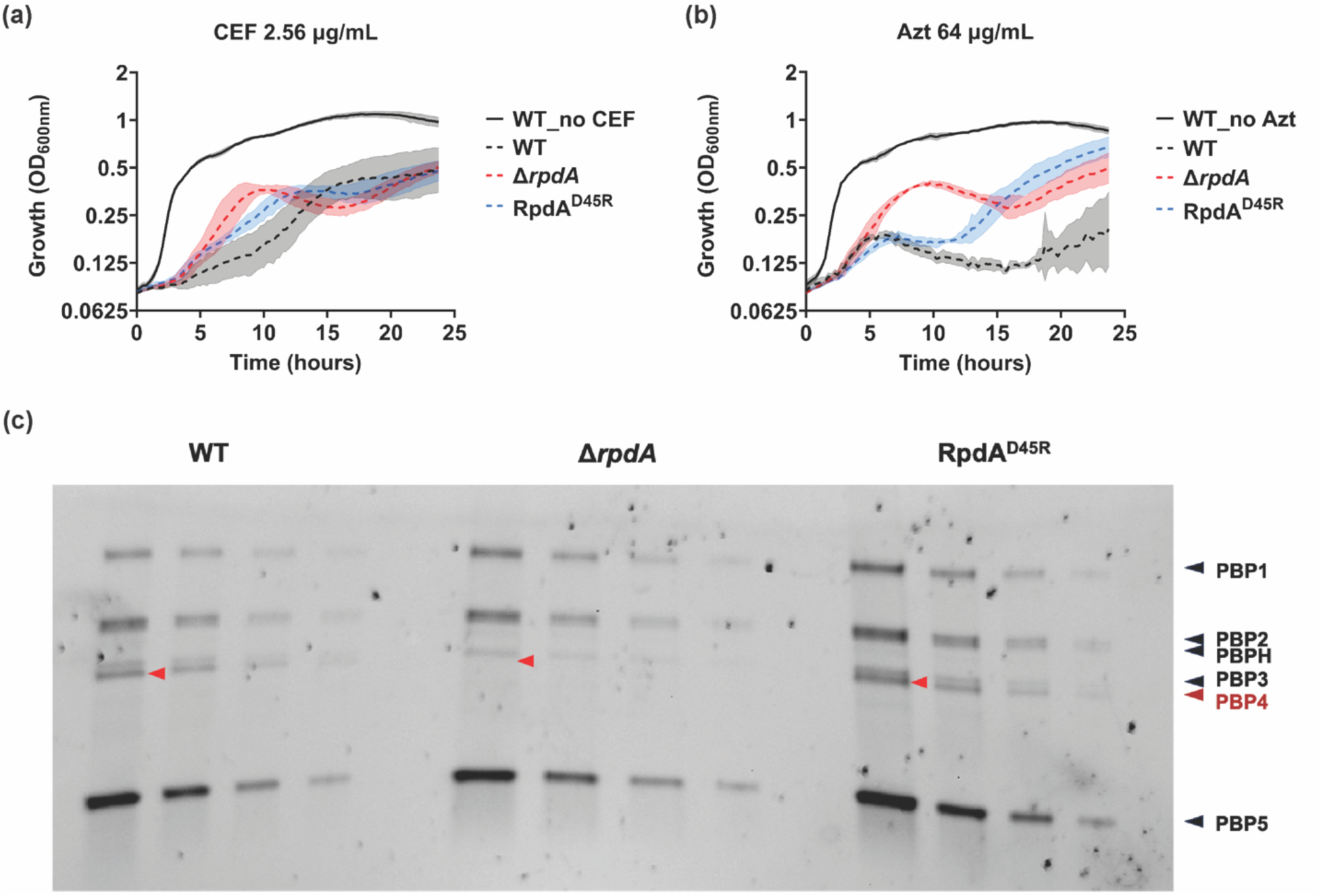
RpdA may have additional role in PBP4 activity. Growth kinetics of WT (black), Δ*rpdA* (red) and RpdA^D45R^ (blue) strains on treatment with **(a)** 2.56 μg/mL CEF and **(b)** 64 μg/mL AZT (dashed lines). Black solid lines show WT growth in the absence of antibiotics. **(c)** PBP4 activity in the membrane fraction of WT, Δ*rpdA* and RpdA^D45R^ strains detected using Boc-FL activity probe. 12, 6, 3 and 1 μL of 0.25 μg/mL of total protein was loaded for each strain (from left to right). Red arrows are used to highlight the loss of PBP4 band in the Δ*rpdA* strain but not in the RpdA^D45R^ mutant. Boc-FL probe assay was performed for 3 biological replicates with two other replicates shown in SI Fig. 8.

### RpdA has a minor role associated with UndP transporter activity

How RpdA may affect PBP4 activity is not yet clear. For example, the extracellular interaction between RpdA and the transglycosylase (TG) domain of PBP4 may increase TG activity, either directly or indirectly. When we compared the membrane topology of RpdA with other transmembrane proteins involved in PG synthesis, the greatest similarity was noted with UptA, which also has 5 transmembrane helices (Fig. 8A). UptA functions as the primary transporter that recycles undecaprenyl phosphate (UndP) back to the cytoplasmic side of the membrane (Roney & Rudner, 2023). Five additional UptA paralogs (COG0586 subfamily DedA proteins) have been identified in *B. subtilis* (Todor Herrera & Gross, 2023), but a strain lacking all six transporters had no significant growth defect (Roney & Rudner, 2023). This suggests that UndP transport may involve other proteins or occur by a protein-independent mechanism.

**Fig 8.**
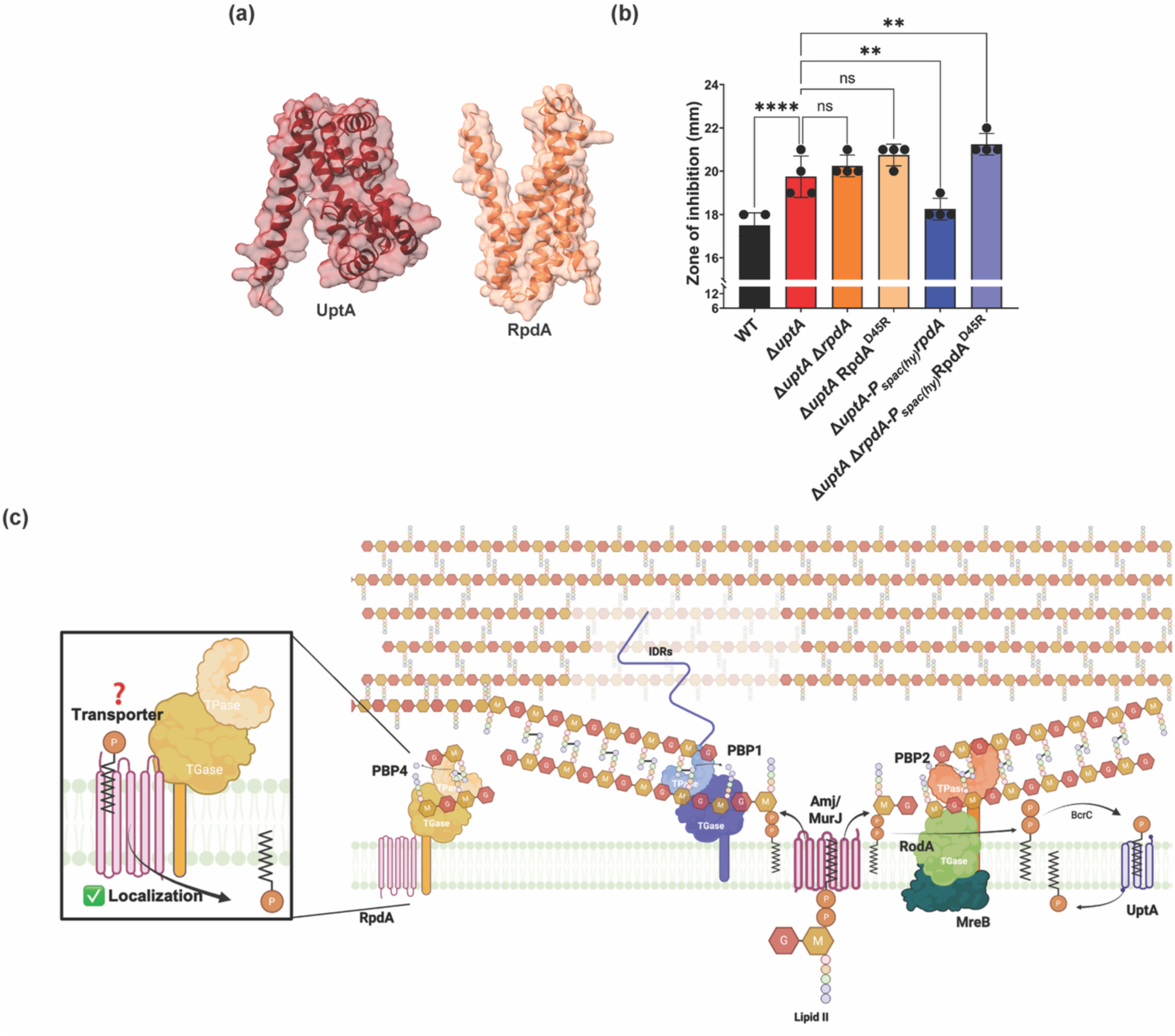
RpdA may have additional roles in supporting PBP4 activity. **(a)** AlphaFold v3 predicted structures of UptA and RpdA. **(b)** Disk diffusion assay of WT, Δ*uptA*, Δ*uptA* Δ*rpdA*, Δ*uptA* RpdA^D45R^ and Δ*uptA* with ectopic expression of *rpdA* and RpdA^D45R^ using the leaky P*_spac(hy)_* promoter under non-inducing conditions. 50 μg MX2401 was loaded on each disk. Data are represented as mean ± SD of at least three biological replicates. Statistical significance was determined by one-way ANOVA with Dunnett’s correction; **** indicates *p* < 0.0001; ns indicates not significant. **(c)** Schematic representation of PG synthesis. PG monomers are synthesized in the cytoplasm and linked to the lipid carrier forming lipid II. Lipid II is flipped to the extracytoplasmic side by MurJ or AmJ. Here, the disaccharide-pentapeptide units are incorporated into the PG meshwork by aPBPs (PBP1 and PBP4) or the elongasome (RodA and PBP2) leaving behind UndPP. UndPP is dephosphorylated by BcrC to generate UndP, which is then recycled back to the cytoplasmic side by UptA. The intrinsically disordered region (IDR) in PBP1 is important for localizing PBP1 to gaps in the PG meshwork, allowing it to repair the PG breaks. The PBP4–RpdA complex has been enlarged for clarity. RpdA functions as a scaffold that localizes PBP4 to the membrane and also provides a minor UndP transport activity. Additional roles for RpdA may exist but were not identified within the scope of this study.

To test the role of RpdA as an UndP transporter, we used MX2401; a derivative of amphomycin that binds to the phosphate group of UndP (Rubinchik *et al*, 2011). Because UptA masks the contribution of all other paralogs, all the strains used in this assay were constructed in a Δ*uptA* background. Using disk diffusion assays, we confirmed that *uptA* deletion significantly sensitized the cells to MX2401 (Fig. 8B). Neither RpdA deletion nor substitution of RpdA with its variant form RpdA^D45R^ affected the sensitivity of the Δ*uptA* strain. However, expression of RpdA with *P_spac(hy)_* significantly reduced the sensitivity of Δ*uptA* (Fig. 8B) Strikingly, induction of the RpdA^D45R^ variant could not reduce the sensitivity of Δ*uptA*. These results support the hypothesis that RpdA functions as a minor UndP transporter and that this activity depends on the interaction between the extracellular helices of RpdA and PBP4.

## Discussion

Biochemical studies in the 1950s established the bacterial cell wall as a single large macromolecule, designated a murein sacculus (Weidel & Pelzer, 1964), comprising glycan strands of alternating NAG-NAM residues crosslinked by short peptide chains (Salton, 1953; Work, 1957). The diverse and often redundant enzymes required for PG synthesis and maintenance are now defined (Anderson *et al*., 2025; Egan Errington & Vollmer, 2020), with several major advances in the last decade (Kumar *et al*., 2022; Zhao *et al*, 2017). This includes the assignment of SEDS family proteins as transglycosylases for the elongasome (RodA) and divisome (FtsW) (Emami *et al*., 2017; Meeske *et al*., 2016) and transporters that flip the lipid II precursor to the exterior face of the membrane (Meeske *et al*, 2015) and recycle the UndP lipid carrier (Roney & Rudner, 2023; Sit *et al*, 2023; Todor Herrera & Gross, 2023). A major remaining challenge is to define the dynamic interactions between enzymes, their complexes, and the sacculus that coordinate PG synthesis, a process involving the localized degradation by autolysins (to allow sacculus expansion), insertion and crosslinking of new glycan strands, and repair of gaps.

PBPs have long been recognized as the major enzymes that assemble the sacculus on the exterior face of the membrane from a lipid-linked, disaccharide-pentapeptide precursor (lipid II) (Sauvage *et al*., 2008). Bifunctional aPBPs have both TG and TP activities and may function in coordination with elongasome and divisome complexes or as solo actors in a diffusive manner. For the latter function, aPBPs are responsible for filling in gaps left by the elongasome (Cho *et al*, 2016; Daitch & Goley, 2020; Vigouroux *et al*, 2020). In *B. subtilis*, the balance between the activity of the elongasome and freely diffusing aPBPs defines the cell shape and width, with elevated activity of the elongasome associated with longer and thinner cells (Dion *et al*., 2019).

Although emerging models suggest that aPBPs may largely function independent of the SEDS-bPBP complexes, their activities are still regulated in ways that are only beginning to be understood. In *E. coli*, the two major aPBPs are both activated by outer membrane lipoproteins, LpoA and LpoB, that penetrate gaps in the sacculus and activates aPBPs (Paradis-Bleau *et al*., 2010; Sardis *et al*, 2021; Typas *et al*, 2010). In *B. subtilis*, a distinct mechanism has evolved to sense gaps in the sacculus: the major aPBP (PBP1) appears to use a carboxy-terminal intrinsically disordered region to identify gaps in the PG matrix (Brunet *et al*., 2022). A variety of other protein-protein interactions are also thought to help regulate PBP activity. For example, in *S. pneumoniae*, PBP1a and PBP2a depend on CozE (Fenton *et al*, 2016) and MacP (Midonet *et al*, 2023), respectively. In *B. subtilis*, EzrA and the GpsB scaffold protein facilitate the septal localization of PBP1 (Cleverley *et al*, 2014; Cleverley *et al*., 2019), and recent findings suggest that cell division in *Clostridioides difficile* requires an aPBP (Harrison & Shen, 2025).

Our study adds RpdA as a new member to this emerging class of partner proteins that help to regulate the activity of PBPs. Our results lead us to propose that RpdA plays two distinct roles: as a scaffold that stabilizes PBP4 in the membrane and as a minor UndP-translocase. Structural predictions revealed a probable interaction between the membrane-spanning α-helices of RpdA and PBP4 that may be crucial for stabilizing PBP4 within the membrane. Consistently, the original loss-of-function mutations we identified in RpdA truncate this segment of the protein (Table S1).

In addition to this membrane-embedded interface, we identified a second predicted contact between an extracellular helix of RpdA and the transglycosylase (TG) domain of PBP4 (Fig. 5). Disruption of this interaction by RpdA^D45R^ mutation, increased CEF and AZT resistance without affecting PBP4 localization (Fig. 7), suggesting that RpdA contributes to PBP4 activity through mechanisms beyond membrane stabilization alone. Using MX2401 sensitivity (Fig. 8B), we further showed that this additional role may involve UndP recycling. Although RpdA is similar in overall size and membrane topology with the known UndP transporter UptA (Roney & Rudner, 2023), RpdA lacks the re-entrant α-helices that are proposed to be important for UndP transport (Todor Herrera & Gross, 2023). This raises the possibility that RpdA may not perform the flipping reaction itself but may instead recruit one of the DedA paralogs to carry out this function. Indeed, we cannot exclude that RpdA recruits other proteins important for PG synthesis such as lipid II flippases or the phosphatases BcrC or UppP that dephosphorylate UndPP to UndP. Further biochemical and structural studies will be required to distinguish between these possibilities and to define the full extent of RpdA’s regulatory roles.

The multiple integrated roles of RpdA in supporting PBP4 activity open a new arena of interactor proteins for aPBPs. Notably, RpdA is conserved across many Firmicutes, and gene-neighborhood analyses show that *rpdA* and *pbpD* are co-located in most species. This conservation and genomic linkage suggest that similar partner–enzyme pairs may be widespread but remain largely unrecognized. Beyond their physiological significance, uncovering such protein partnerships may provide new avenues for antimicrobial target discovery and offer insights into mechanisms of antibiotic resistance.

## Methods

### Reagents and tools table

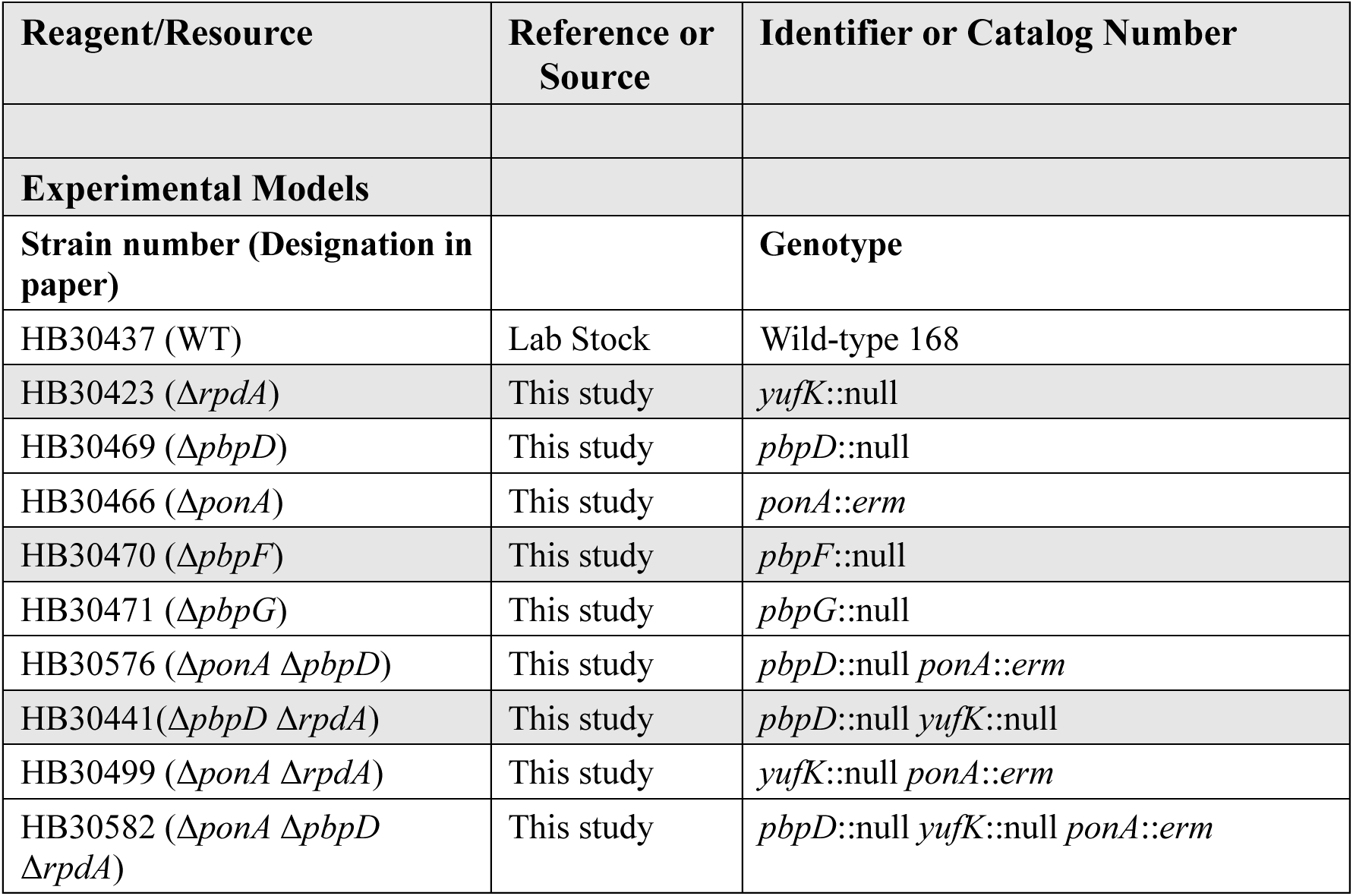
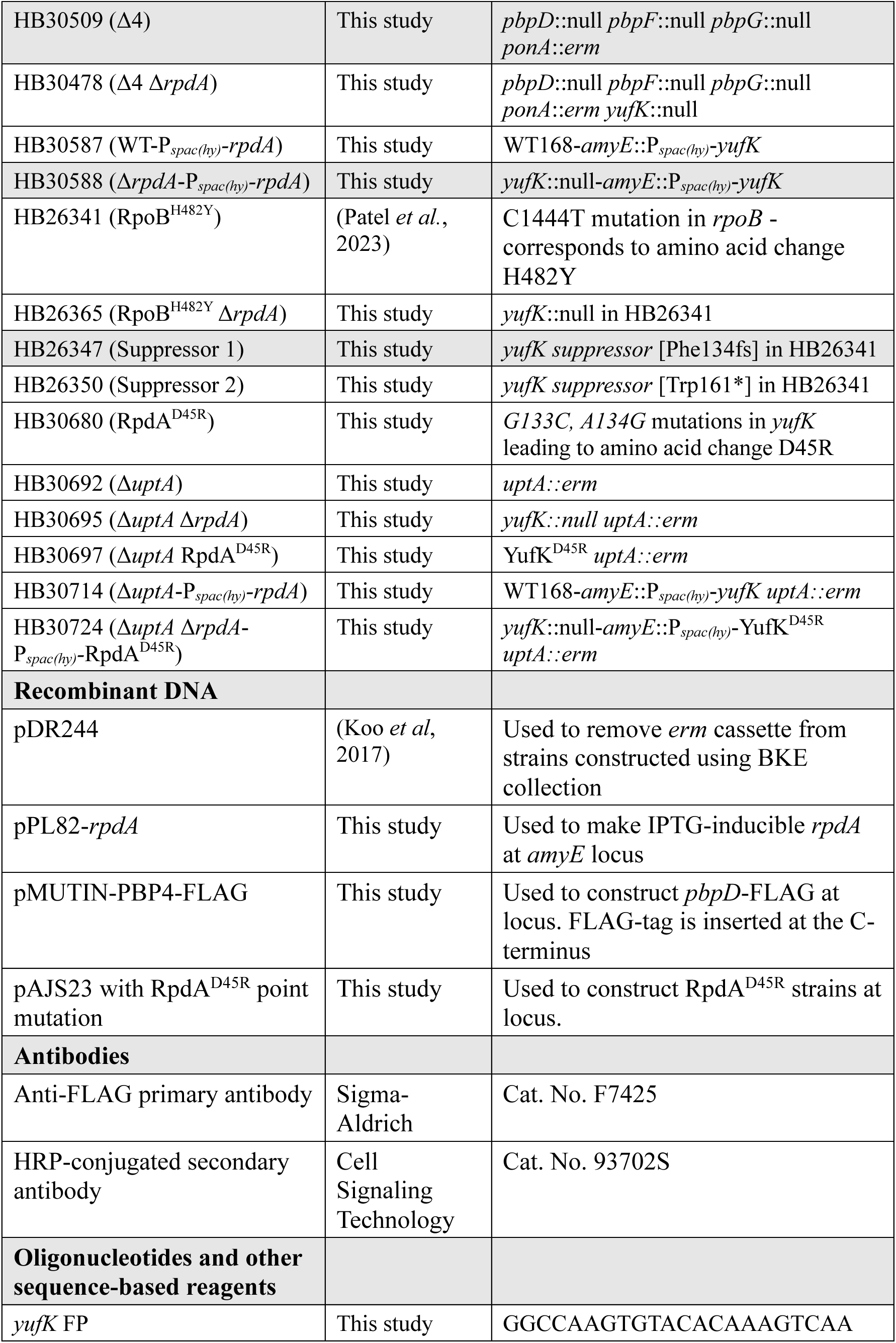
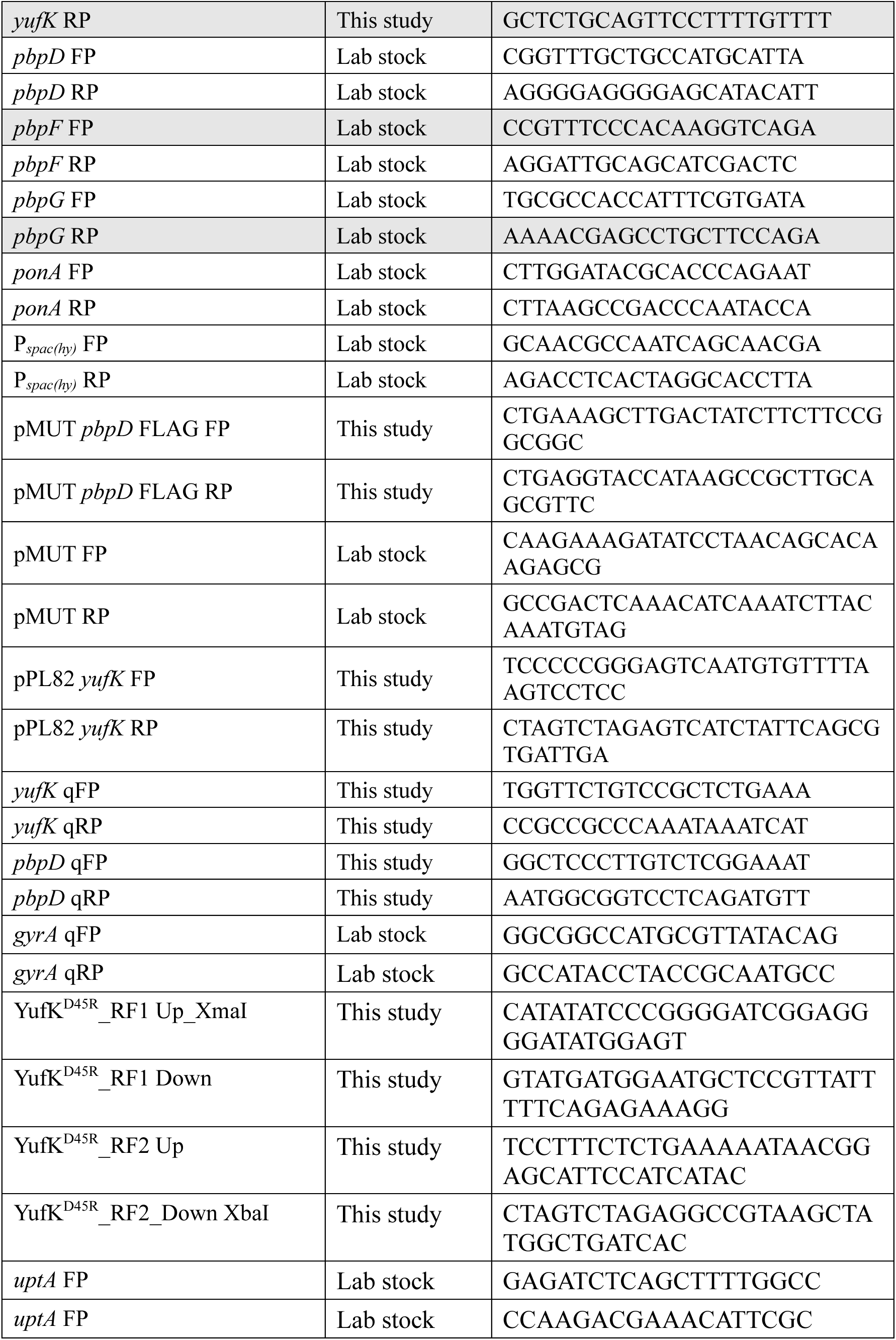
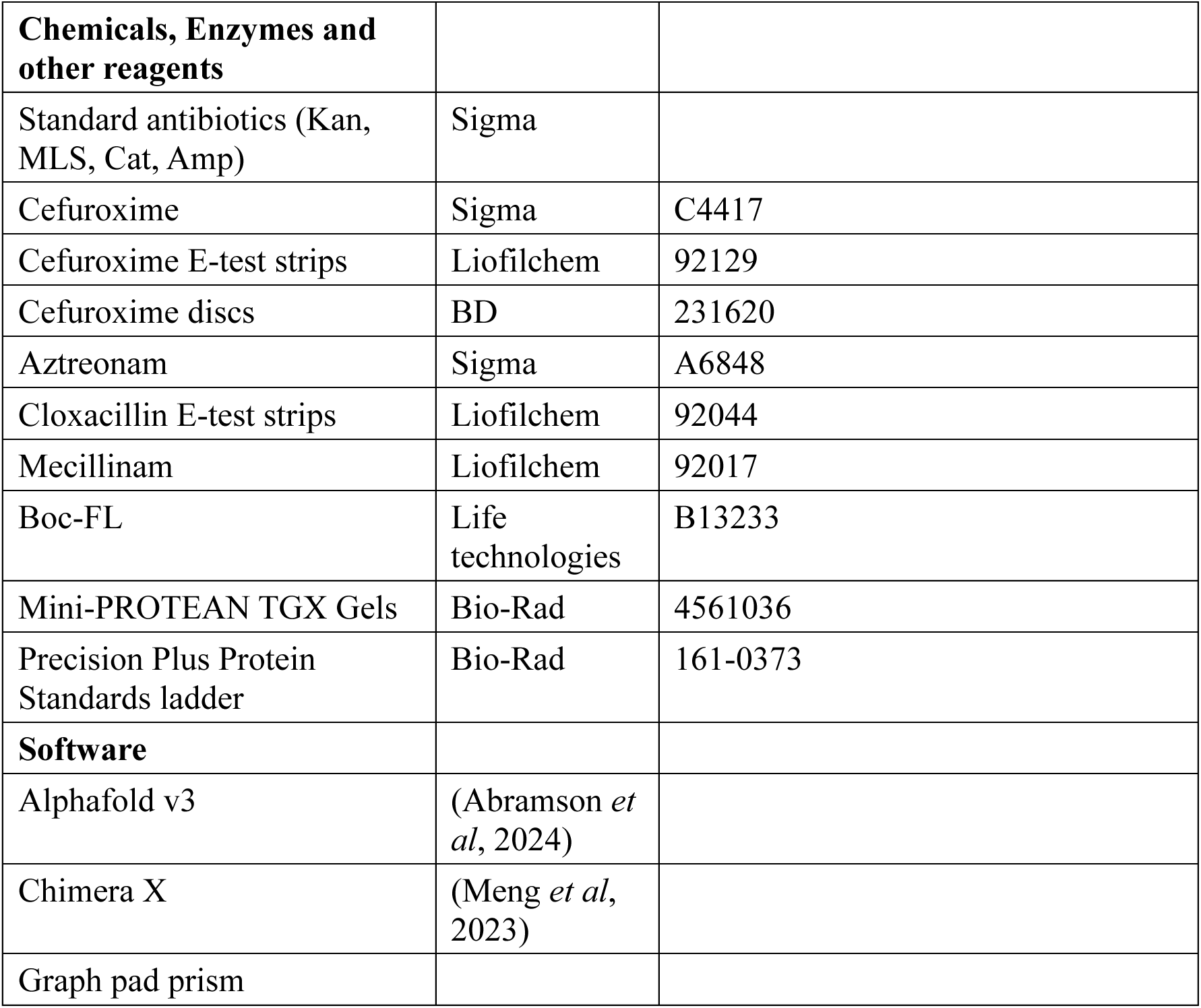

### Bacterial strains

All bacterial strains, plasmids, and oligonucleotides used in this study are listed in the reagents and tools table. Glycerol stocks were streaked out on lysogeny broth (LB) agar plates and incubated overnight at 37°C. For liquid culture, cells were inoculated in LB medium grown at 37°C with shaking at 280 rpm. For transformation, cells were cultured in modified competence medium till optical density at 600 nm (OD_600_) ∼0.8. Gene deletions were constructed using strains from the BKE collection (Koo *et al*., 2017) obtained from National Institute of Genetics, Microbial Physiology Laboratory, NBRP *B. subtilis*. Antibiotics were used at the following final concentrations: 1 μg/mL erythromycin and 25 μg/mL lincomycin in combination for MLS resistance (Macrolide-Lincosamide-Streptogramin B), 10 μg/mL chloramphenicol, 100 μg/mL of spectinomycin. Primers used for strain construction and verification are listed in the reagents and tools table.

The pMUTIN4-FLAG plasmid (Kaltwasser Wiegert & Schumann, 2002) was used to introduce a C-terminal FLAG tag for immunodetection of PBP4. Approximately 500 bp of the 3′ region of *pbpD* was cloned into pMUTIN4-FLAG plasmid, and the FLAG tag was integrated into the chromosome through a single-crossover between the plasmid and the genomic locus. Tagged strains were selected based on MLS resistance conferred by the plasmid and confirmed by Sanger sequencing.

The RpdA^D45R^ strain was constructed at-locus using the pAJS23 CRISPR-Cas9 editing system containing an *erm*-specific guide RNA (Sachla Alfonso & Helmann, 2021). The G133C and A134G mutations in *rpdA* were first introduced by overlap extension PCR to generate the repair template, which was subsequently cloned into pAJS23. The CRISPR editing and selection were performed on plates containing 0.2% mannose and 15 μg/mL kanamycin at 30°C, followed by plasmid curing at 42°C through at least three re-patching steps. Gene-edited strains were identified by the loss of both *erm* and plasmid markers, resulting in MLS- and kanamycin-sensitive phenotypes, and all mutants were verified by Sanger sequencing.

### Disk Diffusion Assay and MIC determination

Antibiotic sensitivity was determined by disk diffusion assay. Bacterial cultures were grown in LB medium at 37°C with shaking to mid-log phase (OD_600_ ∼0.4). A 100 μL aliquot of culture was mixed with 4 mL of molten top agar (0.75% agar) and poured onto 15 mL LB agar plates (1.5% agar). For strains with IPTG-inducible promoters, both the top and bottom agar was supplemented with 0.5 mM IPTG. Once solidified, a 6 mm disk (BD Cat. no. 231620, 30 μg of cefuroxime (CEF) per disk) was placed on the agar. Plates were incubated overnight at 37°C. For this study, the zone of inhibition was defined as the entire region with visibly reduced growth, which includes a central zone with no growth and an outer zone with significantly reduced cell density. For minimum inhibitory concentration (MIC) determinations, Etest strips (Liofilchem Cat No: 92129 (CEF), 92044 (cloxacillin) and 92017 (mecillinam)) were used in place of disks. Plates were incubated at 37°C for 16 hours. MIC values were determined as the point of intersection between bacterial growth and the strip. All experiments were performed with at least three biological replicates. Data are presented as mean ± standard deviation (SD). In histograms, zone of inhibition values were corrected by subtracting the 6 mm disk diameter. Statistical significance was determined using one-way ANOVA with Dunnett’s correction.

### Growth kinetics

200 μL of LB medium with different concentrations of antibiotics was dispensed in each well of a honeycomb microplate. Strains were grown to mid-log phase (OD_600_ ∼0.4) in LB medium, and 1 μL of culture was inoculated into each well. Plates were incubated at 37°C with constant shaking and monitored real-time using Bioscreen Pro analyzer. OD_600_ was measured every 15 min for up to 24 hours. Semi-log growth curves were plotted using GraphPad Prism. Each experiment was independently repeated at least three times. Data are presented as mean ± SD, with the shaded regions indicating SD. Some curves are shown in multiple panels for ease in comparison.

### Plating efficiency

For spot dilution assays, cells were grown in LB medium to mid-log phase (OD_600_ ∼0.4). 25 μL aliquot of culture was diluted into 5 mL of fresh LB medium (1:200) with and without 1.28 μg/mL of CEF and incubated at 37°C with shaking. 1 mL of culture was collected after 12, 15 and 18 hours after treatment. 10-fold serial dilutions were prepared in LB medium and 10 μL of each dilution was spotted on LB agar plates and incubated overnight at 37°C prior to imaging.

### Quantitative real-time PCR (qRT-PCR)

Total RNA was extracted from 1.5 mL of mid-log phase cultures (OD_600_ ∼0.4) cultures using Qiagen RNeasy kit (Cat. No. 74106). Genomic DNA contamination was removed by DNase I treatment (Invitrogen Cat. No. AM1907). RNA concentration and purity were assessed using NanoDrop. For cDNA synthesis, 2 μg of RNA was reverse transcribed using the High-Capacity cDNA Reverse Transcription Kit (Applied Biosystems Cat. No. 4368814). Quantitative PCR was performed using 10 ng of cDNA, SYBR Green Master Mix (Applied Biosystems, Cat. No. 100029284), and gene-specific primers on a QuantStudio 3 Real-Time PCR System. Expression levels of *pbpD* and *rpdA* were measured with *gyrA* as the internal control. Relative expression levels were calculated using the 2^-ΔC^_T_ method and plotted on a log₂ scale. All reactions were performed in technical duplicates, with data obtained from at least three independent biological replicates.

### Bocillin-FL assay

Cells were grown in LB medium at 37°C with shaking till mid-log phase (OD_600_ ∼0.4). 1.4 mL of the cells were harvested by centrifugation at 16,000 *g for 2 min. Cell pellets were washed once with 1 mL of PBS (pH7.4). The same centrifugation settings were used throughout this assay, unless otherwise mentioned. Pellets were resuspended in 50 μL PBS containing 5 μg/mL Boc-FL (0.1% DMSO, Life technologies Cat. No. B13233) and incubated at room temperature for 10 min.

Following labeling, cells were centrifuged and washed once with 1mL PBS. The pellets were then resuspended in 100 μL PBS containing 10 mg/mL lysozyme (from chicken egg white, Sigma Cat. No. L6876) and incubated at 37°C for 30 min. Post incubation samples were sonicated in a BRANSON 1800 water bath sonicator at constant strength for 2 min, followed by 4-6 gentle inversions and a second 2 min sonication. Membrane fractions were isolated by centrifugation at 21,000 *g for 15 min at 4°C and resuspended in 100 μL PBS with 10% glycerol. Protein concentration was determined by Bradford assay, and all samples were normalized to a final concentration of 0.5 mg/mL. 5 μL of each sample was mixed with 15 μL of 2x Laemmli sample buffer, boiled at 90–95°C for 5 min, and then cooled. 12 μL of each sample was loaded onto 10% SDS-PAGE gel (Mini-PROTEAN TGX Gels, Bio-Rad Cat. No. 4561036). The protein bands were separated by electrophoresis for 1 hour and 40 min at 120V. Gels were rinsed three times with distilled water and imaged using a ChemiDoc MP Imaging System (Pro-Q Emerald 488). Each PBP band was identified as reported previously (Sharifzadeh *et al*., 2020), with PBP1 and PBP4 confirmed by the absence of the expected band in the Δ*ponA* and Δ*pbpD* strains, respectively. The experiment was performed with three biological replicates.

### Protein extraction and Western blot analysis

2 mL of overnight preculture was used to inoculate 200 mL LB medium, which was grown until OD_600_ ∼ 0.4. Cells were harvested by centrifugation at 5,000 rpm for 10 min at room temperature. The pellet was washed with 10 mL of TBS buffer (50 mM Tris-HCl, 150 mM NaCl, pH 7.4), then resuspended in 6 mL of Buffer A (TBS, 75 mM sodium azide, 1 mM PMSF) and incubated at 37°C for 60 min. Cells were sonicated in a water bath sonicator at constant power for 5 min, mixed gently 4-6 times by inversion, and sonicated again for 5 min. Supernatant was collected by centrifugation at 12,000 rpm for 15 min at 4 °C. 3 mL of sample was saved as whole-cell lysate, and the rest was fractionated by ultracentrifugation at 100,000 × g for 1 hour at 4 °C. The supernatant was collected as the cytoplasmic protein fraction, while the membrane pellet was resuspended in 1 mL of Buffer B (TBS, 1 mM PMSF). FLAG-tagged proteins were enriched by ANTI-FLAG® M2 Magnetic Beads (Millipore Sigma, Cat. No. M8823). All samples (whole cell-lysates and membrane fraction) were incubated with pre-washed beads for 90 min at room temperature with gentle rotation. Beads were washed three times with TBS buffer, and bound proteins were eluted with 30 μL of TBS. For SDS-PAGE, 30 μL of 2x Laemmli sample buffer containing 5% β-mercaptoethanol was added to each eluted sample, followed by boiling at 95 °C for 5 min. 12 μL of each sample was loaded onto SDS-PAGE gels (Mini-PROTEAN TGX stain-free Gels, Bio-Rad Cat. No. 4568096) along with the Precision Plus Protein Standards ladder (Biorad Cat. No. 161-0373) and transferred to PVDF membrane using a Trans-Blot Turbo Transfer System (Bio-Rad). Membranes were blocked by 5% non-fat dry milk in TTBS (TBS with 0.1% Tween-20) for 1 hour at room temperature. Blots were incubated overnight at room temperature with anti-FLAG primary antibody (1:5000, Sigma-Aldrich Cat. No. F7425), followed by HRP-conjugated secondary antibody (1:5000, Cell Signaling Technology Cat. No. 93702S) for 1 hour at room temperature. Protein bands were visualized using enhanced chemiluminescence (ECL) reagents and imaged with the ChemiDoc MP Imaging System.

### Protein structure analysis

PBP4, RpdA, UptA structures were predicted using AlphaFold v3 (Abramson *et al*., 2024). The protein interaction between PBP4 and RpdA interaction was predicted using AlphaFold v3. The protein complex was visualized and contact residues were predicted using Chimera X.

## Data availability

No large primary datasets were generated or deposited for this study.

## Acknowledgements

We thank Dr. Rudner for providing MX2401. We thank Jessica Willdigg for her contribution during the early stages of this project. We also thank members of Helmann lab for their helpful discussions and technical advice. This research was supported by the National Institutes of Health under award number R35GM122461 to JDH. The content is solely the responsibility of the authors and does not necessarily represent the official views of the National Institutes of Health.

